# Vesicle capture by discrete self-assembled clusters of membrane-bound Munc13

**DOI:** 10.1101/2020.08.17.254821

**Authors:** Feng Li, Venkat Kalyana Sundaram, Alberto T. Gatta, Jeff Coleman, Shyam S. Krishnakumar, Frederic Pincet, James E. Rothman

## Abstract

Munc13 is a large banana-shaped soluble protein that is involved in the regulation of synaptic vesicle docking and fusion. Recent studies suggested that multiple copies of Munc13 form nanoassemblies in active zones of neurons. However, it is not known if such clustering is an inherent self-assembly property of Munc13 or whether Munc13 clusters indirectly by multivalent binding to synaptic vesicles or specific plasma membrane domains at docking sites in the active zone. The functional significance of putative Munc13 clustering is also unknown. Here we report that nano-clustering is an inherent property of Munc13, and is indeed required for vesicle binding to bilayers containing Munc13. Pure Munc13 reconstituted onto supported lipid bilayers assembled into clusters containing from 2 to ∼20 copies as revealed by a combination of quantitative TIRF microscopy and step-wise photobleaching. Surprisingly, only clusters a minimum of 6 copies of Munc13 were capable of efficiently capturing and retaining small unilamellar vesicles. The C-terminal C_2_C domain of Munc13 is not required for Munc13 clustering, but is required for efficient vesicle capture.

## INTRODUCTION

Synaptic vesicle fusion is driven by the zippering of the membrane-proximal helices of v- and t-SNARE proteins Syntaxin1, SNAP25 and VAMP2 into a force-generating bundle termed a SNAREpin (1-3). Syntaxin-1 is initially bound to a dedicated chaperone, Munc18-1, stabilizing a closed conformation that is unable to assemble with SNAP25 and VAMP2 to form a SNAREpin (4). The central MUN domain of Munc13 binds Syntaxin-1 and weakens its interaction with Munc18 (5-8). MUN also independently binds VAMP2 and co-operates with Munc18 to template SNAREpins (9).

Thus, the overall picture that emerges is that only when all three synaptic SNAREs are within molecular contact distance can they be efficiently assembled into SNAREpins, a reaction catalyzed by their two co-operatively acting, specialized molecular chaperones, Munc18 and Munc13 (10). This reaction, further coordinated by the two proteins (Synaptotagmin1 and complexin) that co-operatively clamp the nascent SNAREpins to prevent membrane fusion until signaled by Ca^2+^, primes the vesicles for release, creating what is referred to as the “readily-releasable pool”. Separately but as importantly, Munc13 is important for local capture of the synaptic vesicles, as is needed to bring them within molecular contact range to initiate the cascade of events just outlined (11, 12).

Munc13 was identified from functional screens in nematodes and mammalian synapses (13, 14), and independently as the phorbol ester receptor which mediates drug-enhanced neurotransmitter release (15, 16). Munc13 is a complex protein. Besides its molecule phorbol ester/diglyceride binding C_1_ domain, it also contains three C_2_ domains, and the MUN domain (17). The MUN domain, which contains its chaperone activity is physically about 15 nm in contour “banana”-shaped (18), and is flanked by a C_1_-C_2_ (C_2_B) unit at one end, and a distinct C_2_ (C_2_C) domain at the other end and at the C-terminus of the protein. Its third C_2_ unit (C_2_A) resides at the N-terminus of the protein, and is important for the initial membrane capture of the synaptic vesicle at the active zone, much as the C-terminal C_2_C domain is important for the closer, local capture of the vesicle to trigger SNAREpin assembly (11, 12).

A recent super-resolution optical imaging study of neurons revealed that Munc13 exists in clusters of ∼5-10 copies at the plasma membrane, each cluster associated with a single synaptic vesicle in the readily-releasable pool (19, 20). These results are consistent with the buttressed ring model for vesicle priming, clamping and release (10, 21).

We recently reported results from cryo-electron microscope tomographic reconstructions of docked synaptic-like vesicles in PC12 neuroendocrine cells, which revealed exactly six radially symmetric exocytosis modules under each vesicle (22), each likely containing a single SNAREpin and its associated chaperones, as predicted by the “buttressed rings” hypothesis (10). How exactly 6 SNAREpins can be assembled at each release site is unknown, but is explained in the hypothesis by an outer ring comprised of 6 copies of Munc13, each capable of assembling only one SNAREpin. This outer ring is suggested to enclose an inner ring of the calcium sensor Synaptotagmin-1 which clamps the SNAREpin from fusing. It is envisioned that at an earlier stage Munc13 locally captures the synaptic vesicle in its well-documented erect conformation (10, 23-25) and then re-oriented to hypothetically lie flat on the plasma membrane as it chaperones SNAREpin assembly (with its co-chaperone Munc18) and the assembling SNAREpins forcibly bring the vesicle close to the membrane.

With this background, it was of interest to further investigate in a fully-defined system whether clustering into hexameric or other structures is an inherent property of Munc13, and whether these would be sufficient to capture vesicles.

## RESULTS

### Reconstituted Munc13 molecules form clusters on lipid bilayer membrane

The mammalian (rat) Munc13-1 is a large molecule with 1735 amino acid residues. Here we attached the sequences of a 12x His tag and a Halo to, respectively, the N- and C-termini of its C_1_C_2_BMUNC_2_C domains (residues 529 to 1735, Δ1408-1452, EF, Δ1533-1551 which we called Munc13_L_) and expressed in the Expi293 mammalian cells (26). High quality protein of Munc13 was obtained through Nickel-NTA affinity column purification procedure (Supporting Figure S1). The C-terminal Halo tag allowed us to couple a single fluorophore, Alexa488, conjugated with Halo ligand, to each Munc13_L_ molecule (Supporting Figure S1).

We first made liposomes with DOPC, DOPS, PI(4,5)P2, and DAG (63:25:2:10, mol/mol, see Materials and Methods for details) lipids by extrusion, and then burst the liposomes at the bottom of an Ibidi flow cell (27) with MgCl_2_. The resulting bilayer is uniform and smooth (Supporting Figure S2). 10 nM Munc13_L_-Halo-Alexa488 in a buffer solution containing 50 mM HEPES, 140 mM KCl, 10% glycerol, 1 mM MgCl_2_, and 1 mM DTT was then added on top of the bilayer. After incubation for 60 min, Munc13 molecules that were unbound to the bilayer were removed by extensive washing. The flow cell was then mounted on a TIRF microscope. We observed puncta formed by Munc13_L_-Halo-Alexa488 on the bilayer membrane (Figure 1A). In contrast, when there is no lipid bilayer membrane, very few Munc13 particles were observed at the glass bottom of a bare flow cell (Figure 1B).

**Figure 1.**
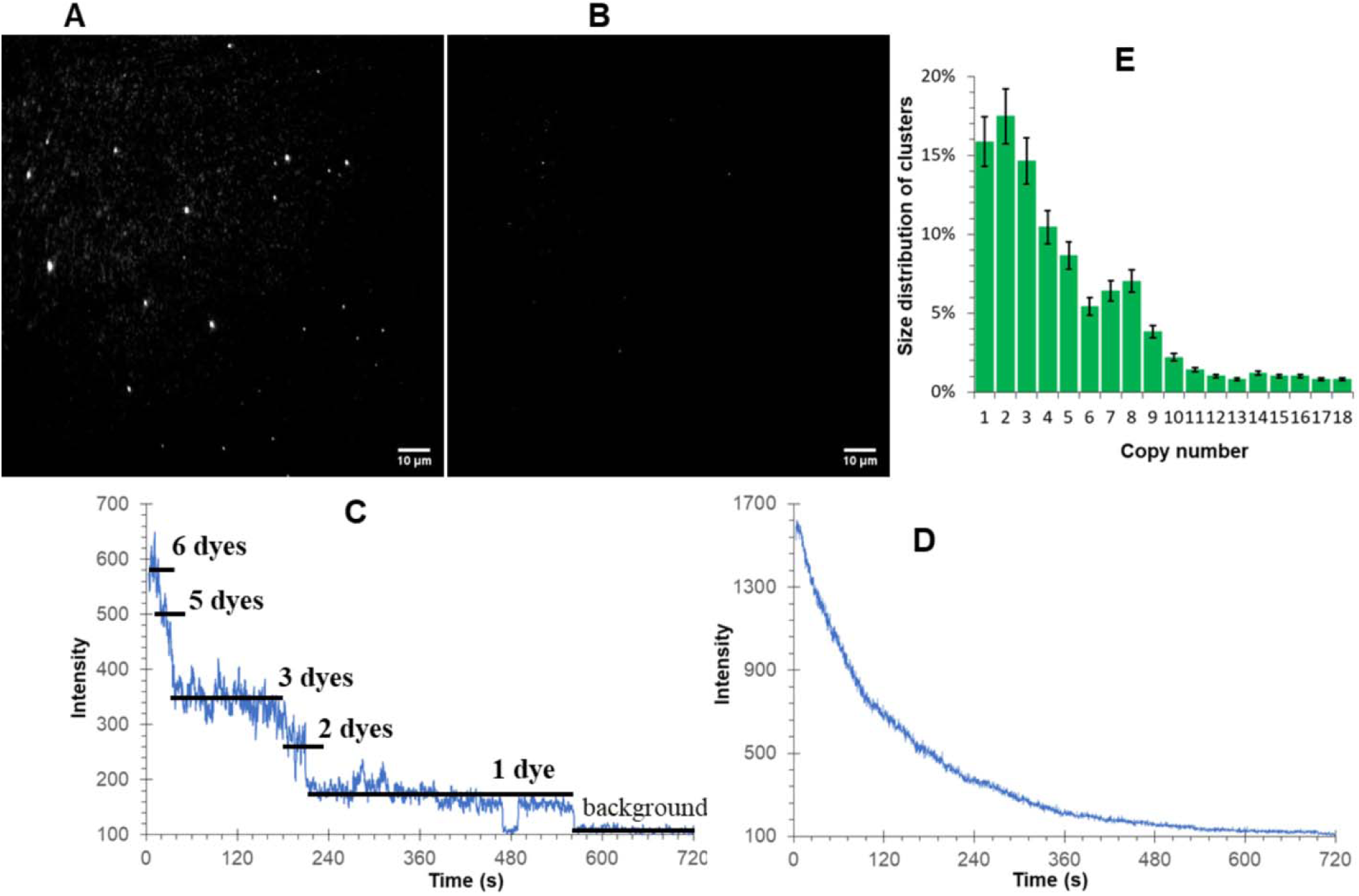
Munc13_L_ form clusters on lipid bilayer. (**A**) & (**B**) Representative TIRF image of particles formed by Munc13 labeled with Alexa 488 on lipid bilayer membrane (**A**) and bare glass surface (**B**). (**C**) & (**D**) Representative step-bleaching profiles of Munc13 clusters of different sizes: small cluster (**C**), where the bleaching steps and corresponding numbers of dyes are indicated; and large cluster (**D**), where the bleaching profile displays a smooth decay. (**E**) Distribution of copy numbers in Munc13 clusters. Error bars represent *s.d*., *n=5*.

As one method with which to determine the number of copies of Munc13_L_-Halo-Alexa488 in each puncta, we gradually bleached the image frames using suitable laser power at different positions (Supporting Movie 1). The bleaching profiles (particle fluorescence intensity versus time) were plotted and a variety of bleaching patterns were found. When the copy number was small, the bleaching profile displayed apparent discrete steps, and the copy number can be determined from counting the number of steps. For example, 6 bleaching steps were found in Figure 1C, each step corresponding to the bleaching of one or two Munc13_L_-Halo-Alexa488 molecules. This method only works for relatively small numbers of copies (generally 5 to 6 or fewer) because the bleaching profile becomes smooth when the copy number is large (Figure 1D). As an alternative for larger copy numbers, we fitted the intensity profile using

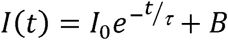

where *I*(*t*) is the intensity at time *t, I*_0_ is the initial intensity before bleaching, *τ* is the time constant, and B is the background (Supporting Figure S3). Hence the copy number N can be obtained through

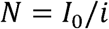

where *i* is the unit intensity of a single fluorophore or the average size of bleaching steps, such as in Figure 1D.

The histogram of the copy number shows that the size of the clusters has a broad distribution from 1 to 18 molecules (Figure 1E) with a large fraction of smaller oligomers. The average density of clusters is about 5 ± 1 clusters per 1000 μm^2^. Therefore the average distance between two nearest neighboring clusters is 14 ± 2 μm. This suggests that the Munc13 clusters are well separated from each other and must form independently.

In principle there are two possible ways Munc13 clusters could form: (i) the clusters could self-assemble via direct contacts between the Munc13 molecules themselves; or (ii) Munc13 molecules could independently bind to a common sub-resolution (<250 nm) lipid domain enriched for example in the acidic lipids PIP_2_ and/or PS,(28) and simply appear to be molecularly clustered. In the first model Munc13 may be able to cluster when free in solution; in the second model this could not occur.

To test this (Fig. 2) we examined the state of assembly of Munc13_L_ in solution, applying samples directly to a cover slip and examining individual molecules of Munc13_L_-Halo-Alexa488, having been previously incubated either in the absence of phorbol ester or in its presence (250 nM). Phorbol esters were reported to trigger relocation of Munc13 from cytosol to the plasma membrane in cells.(15) Clusters of Munc13_L_ were indeed observed by TIRF in the presence (Figure 2A) but not in the absence of phorbol ester (Figure 2B). With phorbol ester, the size distribution of the clusters is similar to those formed on lipid bilayers (Figure 2C). In contrast, in the absence of soluble phorbol ester, Munc13_L_ exists mainly as monomers and dimers. Particles settled on glass surface and displayed a more uniform distribution (Figure 2D). Thus Munc13_L_ clusters direct self-assembly mechanism.

**Figure 2.**
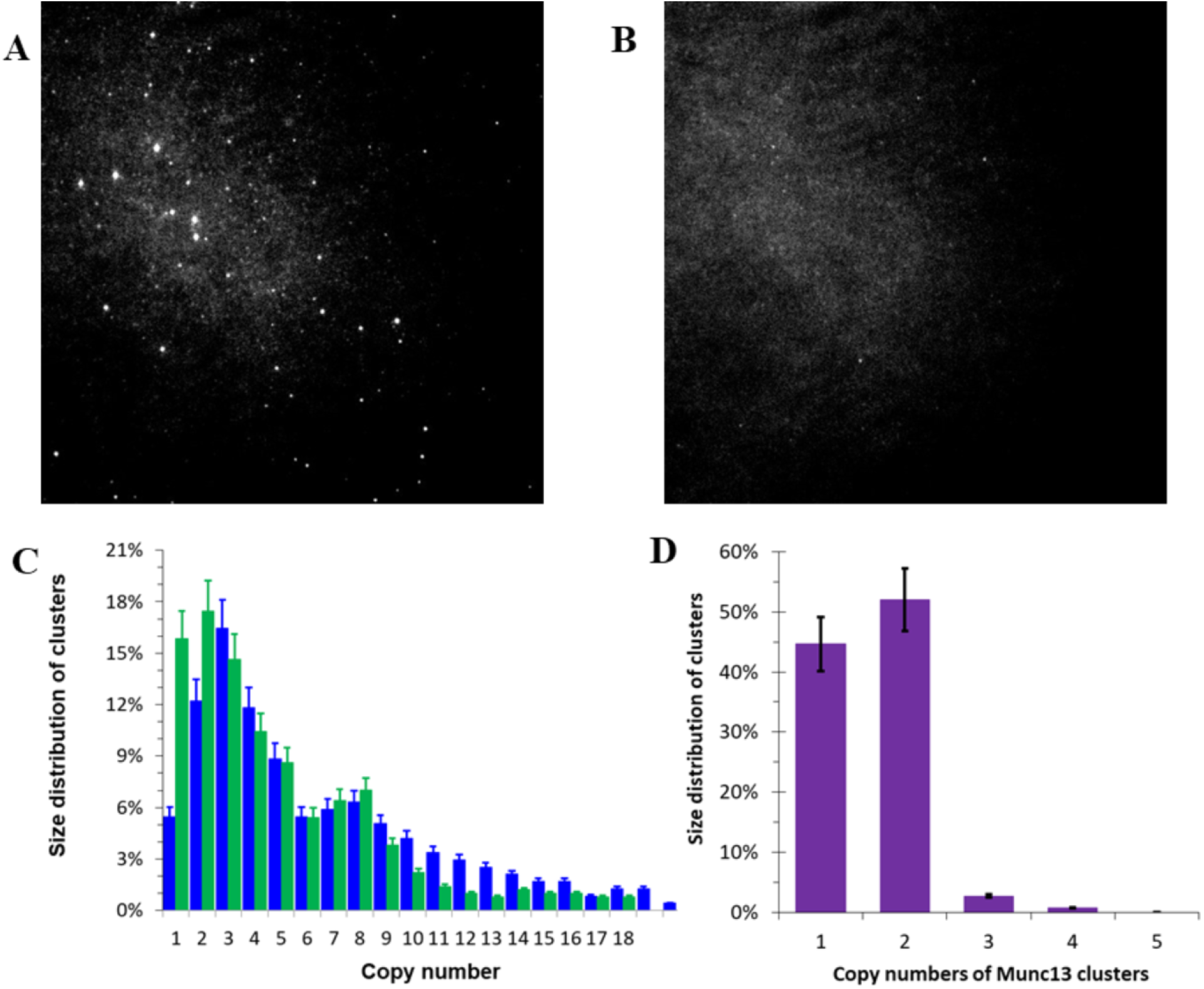
Munc13_L_ form clusters in solution with the presence of phorbol ester. (**A**) & (**B**) Representative TIRF image of particles formed by Munc13 labeled with Alexa 488 in solution with 250 nM phorbol ester (**A**) and without phorbol ester (**B**). **(C**) Size distribution of Munc13 clusters formed in solution with 250 nM phorbol ester (blue columns). The green columns are the size distribution of Munc13 clusters formed on lipid bilayer membrane, which serves as a reference. (D) Distribution of copy numbers in Munc13 particles formed in solution in the absence of phorbol ester.

### Munc13_L_ clusters stably capture vesicles to lipid bilayer membranes

Next we tested the ability of Munc13_L_ clusters to stably capture vesicles. We incubated the Munc13_L_-bound bilayer with vesicles that contained DOPC, DOPS, DPPE-Atto647N (68:30:2, mol/mol, see Materials and Methods for details), followed by an extensive buffer wash, so that only vesicles that were stably bound to the membrane would remain. Clusters of Munc13_L_ and captured vesicles were independently imaged in the 488 nm and 633 nm channels, respectively (Figure 3A). The average density of vesicles on lipid bilayer membrane was about 2.5 ± 0.5 vesicles per 1000 μm^2^.

**Figure 3.**
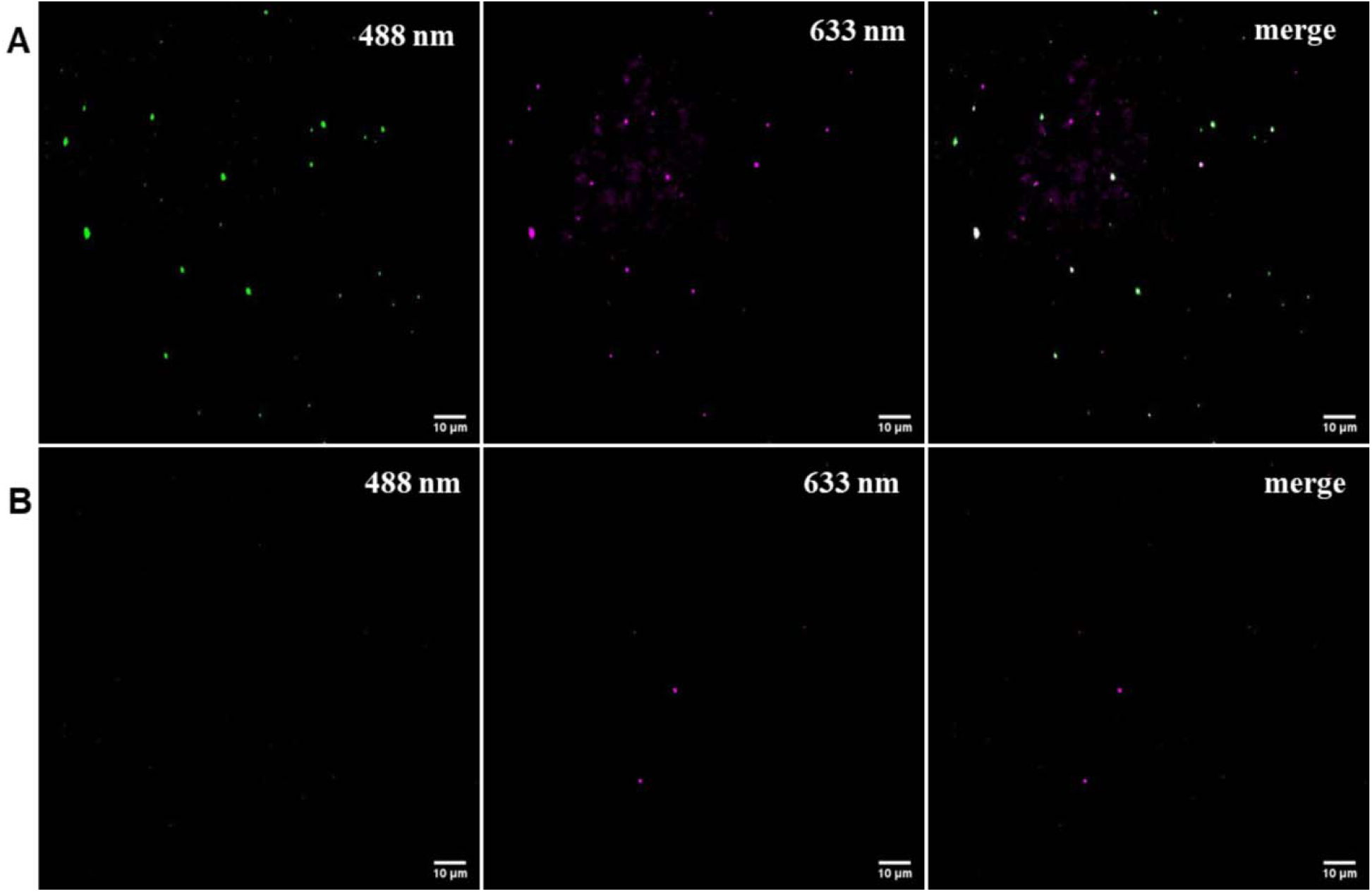
Munc13_L_ clusters capture vesicles to lipid bilayer. (**A**) Munc13 clusters co-localize with vesicles which are captured to lipid bilayer from solution: TIRF image of clusters formed by Munc13_L_ labeled with Alexa 488 on lipid bilayer (*left panel*), and TIRF image of vesicles containing DOPE-Atto647 lipid that are anchored to lipid bilayer (*middle panel*), and merge of the previous two images (*right panel*). About ∽75% vesicles on the lipid membrane are bound to Munc13 particles. (**B**) In the absence of Munc13, no specific capture of vesicles to lipid bilayer: TIRF image of the Alexa 488 channel (*left panel*), and TIRF image of vesicle containing DOPE-Atto647 lipid that are anchored to lipid bilayer (*middle panel*), and merge of the previous two images (*right panel*).

Merging the images from these two channels revealed that about 75 ± 15% of all the membrane-attached vesicles were localized to Munc13_L_-containing clusters (Figure 3A). As the control experiment, we incubated vesicles with the bilayer membrane in the absence of Munc13 (Figure 3B). The average density of vesicles on lipid bilayer membrane dramatically decreased to ∽ 0.5 vesicles per 1000 μm^2^, which was similar to density of vesicles that were not co-localized with Munc13 in Figure 3A, suggesting they were non-specifically attached to the bilayer.

Thus, Munc13_L_ clusters are inherently able to capture vesicles from solution and stably tether them to the bilayer membrane. Previous studies related to Munc13 capture of vesicles have either been in vivo, showing that Munc13 is necessary for vesicle recruitment but not informing on whether Munc13 acts directly or is sufficient (12); or they have been in defined systems showing clustering of homogeneous vesicles (11, 17), a process that is not necessarily related to vesicle capture of heterotypic vesicles by plasma membrane.

### Munc13_L_ clusters must be at least hexamers to capture vesicles

Not all the Munc13_L_ clusters captured vesicles in the previous experiments (Figure 3A). To establish whether there is relationship between the size of a Munc13_L_ cluster and its capacity for vesicle capture, we determined the copy number of each cluster on the image frame from the step-bleaching profiles and recorded whether or not it had captured a vesicle (Figure 4A). We took the data sets from 5 image frames drawn from independent experiments. Figure 4B reveals that the copy number distributions of Munc13_L_ clusters that have captured a vesicle (red) and those that have not (blue) are markedly different. A distinct transition was observed: most clusters that did not capture a vesicle had 5 or fewer copies of Munc13_L_, whereas those clusters that had stably captured a vesicle contained 6 or more copies.

**Figure 4.**
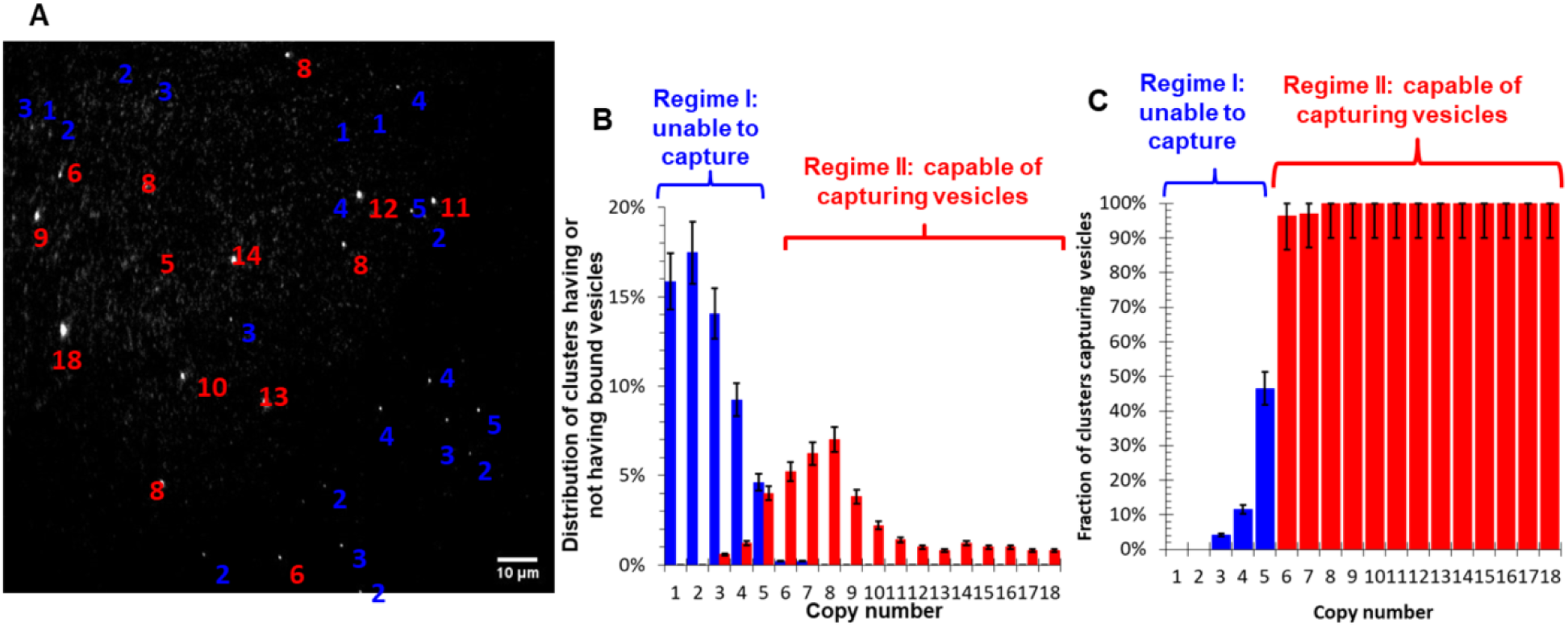
The copy number in Munc13_L_ clusters needs to exceed a threshold value to be capable of capturing vesicles. (**A**) Mapping of copy numbers of Munc13_L_ clusters in a representative image frame and these clusters’ capability of capturing vesicles. The numbers designate copy number of the clusters, and red color refers to clusters that are capable of capturing vesicles, whereas blue color refers to clusters that incapable of capturing. (**B**) Correlation between copy numbers of Munc13_L_ clusters and their capability of capturing vesicles. Copy number distribution of Munc13_L_ clusters that are capable (red) and incapable (blue) of capturing vesicle, respectively. (**C**) Probability of capturing one or more vesicles by Munc13_L_ clusters with different copy numbers.

Drawing on the same data set, we instead calculated the probability that a cluster captured a vesicle as a function of the number of Munc13_L_ molecules in the cluster (Figure 4C). This analysis revealed that clusters has 6 or more copies of Munc13 had a nearly 100% probability of stably capturing a vesicle, and this probability becomes insignificant when there are fewer than 5 copies.

We conclude that a hexameric cluster contains the minimum, threshold number of copies of Munc13_L_ needed to reliably and stably capture a single vesicle in a fully-defined system.

### The C_2_C domain is important for vesicle capture but not for clustering

The C-terminal C_2_ domain of Munc13, C_2_C domain, was reported to be important in bridging liposome vesicle membranes (11). To test whether this domain affects the formation of Munc13 clusters and their vesicle ability to capture vesicles, we generated a construct in which the C_2_C domain was deleted, Munc13_S_. Like the Munc13_L_-Halo construct with C_2_C, a Halo tag was added on the C-terminus for subsequent labeling. This C_1_C_2_BMUN-Halo construct, amino acid residues 529 to 1531 (Δ1408-1452, EF), is well expressed in Expi293 cells. After purification using the Nickel-NTA column, the protein displayed a single band on the SDS-PAGE gel.Purified protein was then labeled using the Halo ligand that was conjugated with Alexa488, so that a single fluorophore was coupled to every ΔC_2_C Munc13_S_ molecule (Supporting Figure S1).

As previously, we then prepared bilayers that contained PC, PS, PIP2, and DAG lipids. As a control, we used a bare flow cell. Both the bilayer and control cases were incubated with ΔC_2_C Munc13. After washing away the unbound protein, Munc13 puncta formed by ΔC_2_C Munc13_S_-Halo-Alexa488 on the bilayer membrane were observed (Figure 5A). These particles displayed comparable features as the clusters formed by Munc13_L_ containing C_2_C domain.

**Figure 5.**
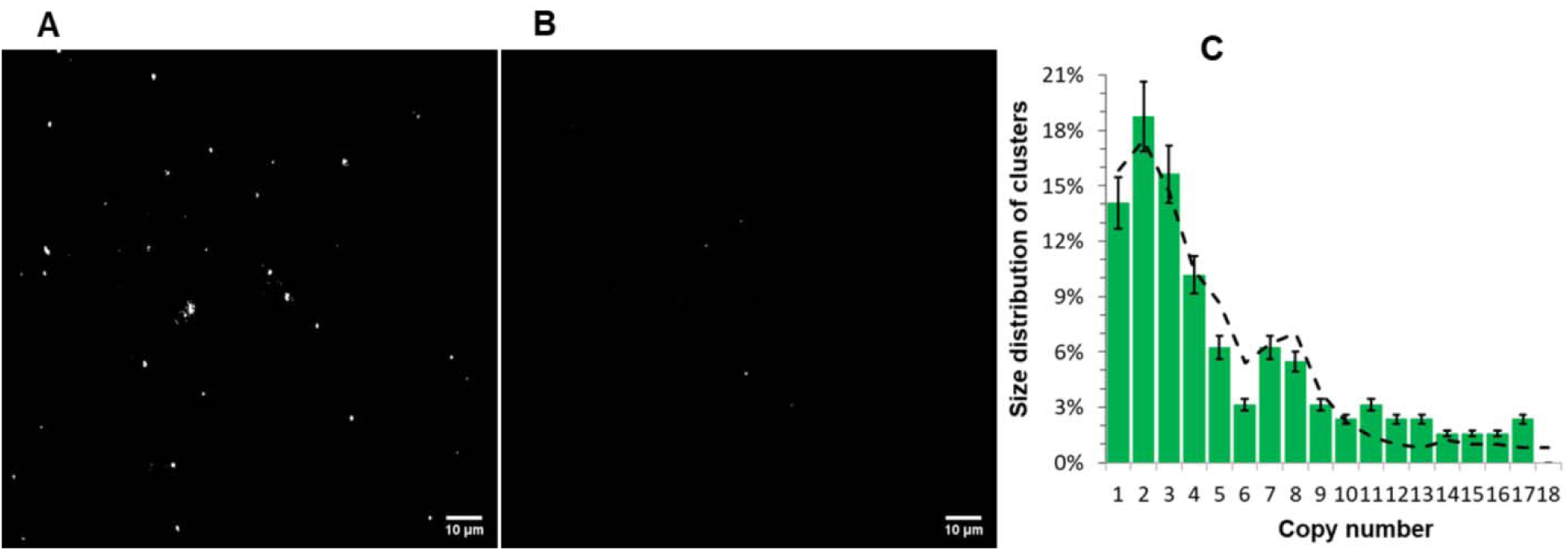
C^2^C domain does not affect Munc13 clustering. (A) & (B) Representative TIRF image of particles formed by Munc13_S_ ΔC^2^C labeled with Alexa 488 on lipid bilayer membrane (A) and bare glass surface (B). (C) Distribution of copy numbers in Munc13_S_ ΔC^2^C clusters. The dashed line is the size distribution of Munc13_L_ clusters with C^2^C domain, which serves as a reference.

First, when there is no lipid bilayer membrane, very few ΔC_2_C Munc13_S_ particles were observed on bare glass surface (Figure 5B). This suggested that ΔC_2_C Munc13_S_ did not form clusters in the absence of lipid bilayer. Moreover, the ΔC_2_C Munc13_S_ formed clusters on lipid bilayer.When puncta of ΔC_2_C Munc13_S_ on bilayers were gradually bleached using the appropriate laser power, as before a mixture of large clusters with high initial intensity and continuous and smooth bleaching profiles, and small clusters with lower initial intensity could be observed (Supporting Movie 2). Using the same analytical methods as before, we found that the size distribution of the ΔC_2_C Munc13_S_ clusters was similar to the distributions of Munc13_L_ clusters containing the C_2_C domain, (Figure 5C). This result establishes that the C_2_C domain is not involved in the self-assembly process that results in the observed clustering of Munc13.

To test for vesicle capture, the bilayer sample was incubated with vesicles that contained DOPC, DOPS, DOPE-Atto647N (68:30:2, mol/mol). Despite the similarity in size distribution of Munc13 clusters with and without C_2_C domain, the image frames of 633 nm channel showed that the number of vesicles bound to ΔC_2_C Munc13_S_ bilayer was markedly decreased (Figure 6A-C).Finally, we correlated the size of ΔC_2_C Munc13_S_ clusters with their ability to capture vesicles. The fraction of ΔC_2_C Munc13_S_ clusters that captured vesicles was greatly decreased as a result of deletion of C_2_C, and now even clusters of 6 or more copies failed to capture them reliably (Figure 6D).

**Figure 6.**
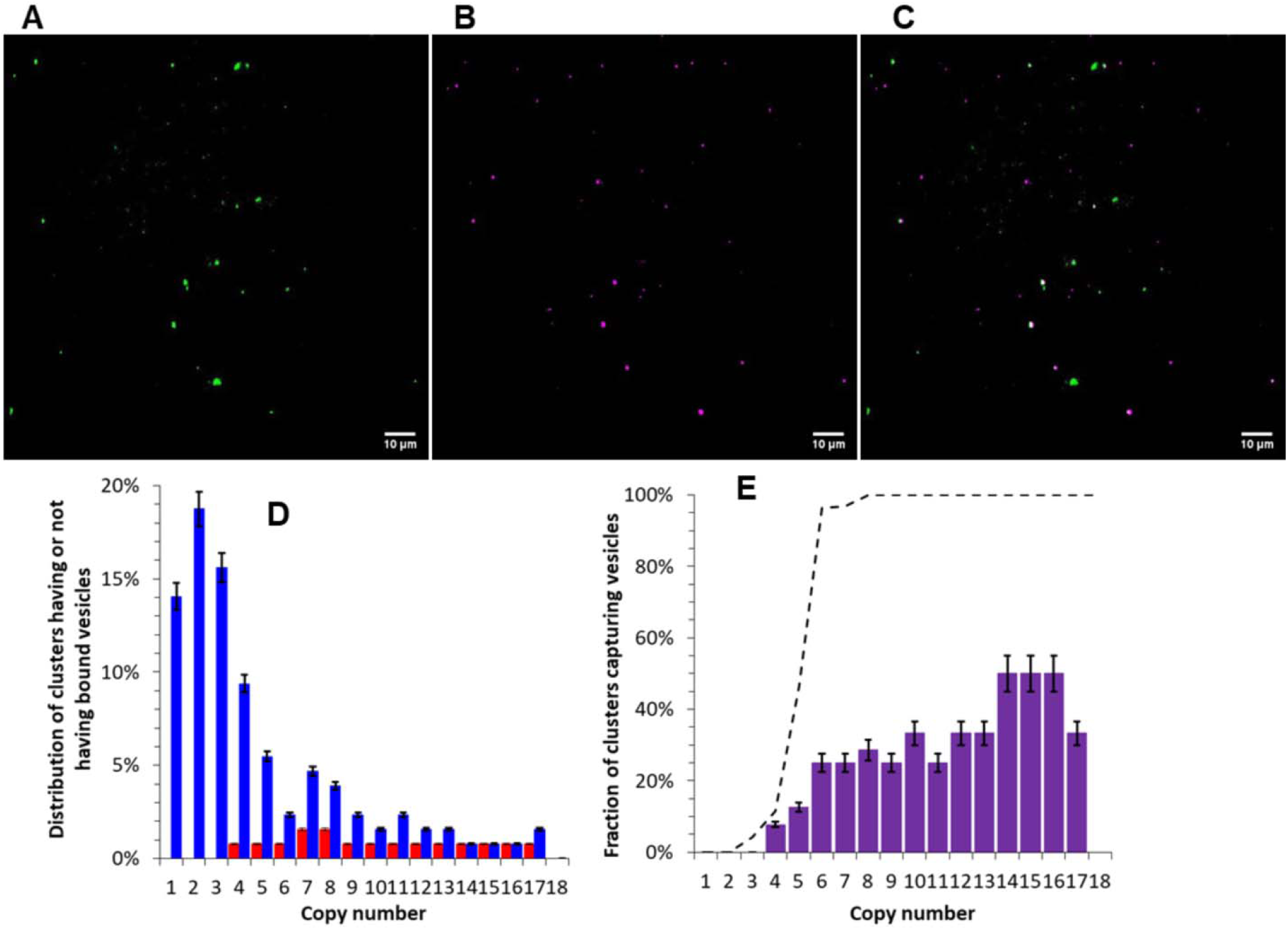
C^2^C domain affects Munc13’s function of capturing vesicles. (**A**) TIRF image of clusters formed by Munc13_S_ ΔC^2^C labeled with Alexa 488 on lipid bilayer. (**B**) TIRF image of vesicle that are anchored to lipid bilayer. (**C**) Merge of the previous two images, which shows that Munc13_S_ clusters capture fewer vesicles in the absence of C^2^C domain. About ∽31% vesicles on the lipid membrane are bound to Munc13 particles. (**D**) Copy number distribution of Munc13_S_ ΔC^2^C clusters that are capable (red) and incapable (blue) of capturing vesicle, respectively. (**E**) Probability of capturing vesicles by Munc13_S_ ΔC^2^C clusters with different copy numbers. The dashed line is the same probability by Munc13 clusters with C^2^C domain, which serves as a reference.

With C_2_C domain, Munc13 clusters had nearly 100% probability of capturing vesicles when their copy numbers were 6 or more. However, for ΔC_2_C Munc13_S_ clusters, the likelihood of vesicles capture was reduced by more than half even when they had 6 or more copies of ΔC_2_C Munc13_S_ molecules (Figure 6E). This demonstrates that the C_2_C domain of Munc13 plays a significant role in vesicle capture, possibility by directly interacting with the vesicle membrane. Overall, our results show that cluster formation does not require the C_2_C domain, but deletion of C_2_C significantly affects Munc13’s function of interaction with vesicles.

## DISCUSSION

In this study we reconstituted Munc13 on lipid bilayer membranes, and observed that the protein forms clusters containing 2 – 18 copies, determined from the step-bleaching of individual fluorescent dyes or from the initial fluorescent intensity of the cluster bleaching profiles. Clustering required binding to the bilayer and is an intrinsic property of the pure protein. Because we did not use super-resolution imaging we cannot formally prove that the clusters exist on the nano-scale, as distinct from consisting of individual, unassociated Munc13 molecules that we cannot optically resolve from each other. However, the fact that a unit of at least 6 such copies, clustered *before vesicle binding* within a single diffraction-limited region of bilayer surface, can cooperate to bind a common vesicle places these six copies no further apart than the diameter of an ∼50-100 nm small unilamellar vesicle. Moreover, Munc13 is able to form similar size clusters as a pure protein in solution when triggered to do so by the diglyceride analogue phorbol ester, suggesting that formation of clusters is through direct contacts among Munc13 molecules rather than separate binding to a common lipid domain without molecular contacts among them.

Even though binding to other synaptic proteins such as Syntaxin, Synaptotagmin, or other synaptic proteins is evidently not required for Munc13 clustering, it is entirely reasonable to imagine that such interactions may occur, and could affect the degree of Munc13 oligomerization, imposing constraints that could limit its extent and perhaps shape uniform clusters. For example, in the buttressed ring model an interaction with the inner ring of Synaptotagmin would template the Munc13 oligomer into uniform rings of six copies (10). There are various ways Munc13 molecules could self-assemble into oligomers, either laterally side-to-side or end-to-end, or a combination. It is unlikely that the C_2_C domain is involved because in its absence Munc13 still forms clusters with size distribution similar to the molecules with C_2_C domains (Figure 5).

At presynaptic terminals, vesicles are drawn from reserve and recycling pools for capture at active zones to form the readily releasable pool that is primed for synchronous release. The SNAREs and Synaptotagmin-1 are known to bridge vesicles to the plasma membrane through membrane-insertion (SNAREpins) or binding (Synaptotagmin-1) to PIP2 with strong supporting *in vivo* evidence (28-34). Munc13 similarly is a strong candidate for a vesicle tether based on *in vitro* (11, 17, 35) and *in vivo* studies.(11, 12, 36, 37) Our finding that hexameric clusters of Munc13 are minimally needed to reliably capture vesicles compares favorably with ∼5-10 copies that are anatomically associated with each synaptic vesicle in the readily-releasable pool (19, 20). Our finding that the C_2_C domain is required for reliable vesicle capture is also consistent with *in vivo* data and electron tomography results (11). Therefore, our data support the prevailing model that the Munc13 C_1_C_2_B domains bind to the plasma membrane, with the MUN domain rising up essentially perpendicular to the plasma membrane elevating and presenting the C_2_C domain to capture synaptic vesicles (11, 35).

This model is attractive because it elegantly explains the sequential hierarchy of physical limitations on vesicle-membrane separation at which initial capture can take place due to the different sizes of Synaptotagmin-1, SNAREs and Munc13. The interaction range of Munc13 is about 20 nm (18), whereas Synaptotagmin-1 can first interact with acidic lipids on plasma membrane at about 5 nm (21, 24). Zippering of the SNAREs is triggered when intermembrane distance is about 8 nm (30, 38). Assuming that the capture of vesicles that we have observed *in vitro* is mediated by a perpendicular arrangement of Munc13 in the clusters, this interaction will necessarily precede vesicle attachment by either Synaptotagmin-1 and by SNAREs. If this is correct, nascent SNAREpins will have the opportunity to bind Synaptotagmins and Complexin (39) even before Synaptotagmin can bind the plasma membrane and oligomerize into the proposed inner ring in the buttressed rings model (10).

The surprising requirement that Munc13 clusters have to reach a critical copy number for vesicle capture is currently unexplained. Clusters with of 4 or fewer Munc13 proteins can barely capture vesicles, whereas clusters of 6 or more have capture vesicles with nearly perfect reliability. One possible explanation is that a single C_2_C domain lacks sufficient affinity to reliably capture and stably retain a vesicle, and multivalent binding is required. This seems unlikely because multivalent interactions generally increase smoothly (and geometrically) with the degree of valency. The alternative is that the clusters present a specific architecture needed for binding that cannot form with less than a quorum of 6 subunits. This could be a hexameric arrangement of subunits in the known perpendicular (erect) topology, or it could be a co-planar hexameric ring of Mun13 on the surface of the bilayer, as proposed in the buttressed ring hypothesis, perhaps following a transition from initial capture in the perpendicular state.

## MATERIALS AND METHODS

### Chemicals

The lipids used in this study, 1,2-dioleoyl-sn-glycero-3-phosphocholine (DOPC), 1-palmitoyl-2-oleoyl-sn-glycero-3-phosphocholine (POPC), 1,2-dioleoyl-sn-glycero-3-(phospho-L-serine) (sodium salt) (DOPS), L-α-phosphatidylinositol-4,5-bisphosphate (Brain, Porcine) (ammonium salt) (brain PI(4,5)P2), 1-2-dioleoyl-sn-glycerol (DAG), and 1,2-dipalmitoyl-sn-glycero-3-phosphoethanolamine-N-(7-nitro-2-1,3-benzoxadiazol-4-yl) (ammonium salt) (DPPE-NBD) were purchased from Avanti Polar Lipids. 1,2-Dioleoyl-sn-gylcero-3-phosphoethanolamine ATTO 647N (DOPE-Atto647N) was from ATTO-Tec. 4-(2-Hydroxyethyl)piperazine-1-ethanesulfonic acid (HEPES), Potassium hydroxide (KOH), Potassium chloride (KCl), Magnesium chloride (MgCl_2_), Glycerol, DNAse I, RNAse A, Benzonase, Roche complete protease inhibitor cocktail tablets, Phorbol 12,13-dibutyrate, Phorbol 12-myristate 13-acetate, and DL-Dithiothreitol (DTT) were purchased from Sigma-Aldrich. Nickel-NTA agarose, TCEP-HCl, and Expi293™ Expression System Kit were from Thermo Fisher Scientific. Plasmid maxi prep kit was from QIAGEN. HaloTag® Alexa Fluor® 488 Ligand was from Promega. All aqueous solutions were prepared using 18.2 MΩ ultra-pure water (purified with the Millipore MilliQ system).

### Protein constructs, expression and purification

The original vector expressing rat Munc13 was a kind gift from Dr. Claudio Giraudo. The plasmid expressing the Halo tag (pFN21A) was purchased from Promega. The expression plasmids His_12__PreScission _C_1__C_2_B_MUN_C_2_C_tev_Halo and His_12__PreScission _C_1__C_2_B_MUN_tev_Halo (ΔC_2_C Munc13) were produced by cloning rat Munc13-1 residues 529 to 1735 and 529 to 1531, respectively, to a mammalian cell expression vector between the BamHI and NotI sites. Munc13-1 residues 1408-1452 were deleted and residues EF were added in (5). A short linker sequence containing a TEV cut site was subcloned in followed by the Halo tag. The resulting plasmids were amplified with maxi prep using QIAGEN Plasmid Maxi kit and were used to transfect Expi293F™ human cells. Proteins were expressed with Expi293™ expression system following manufacturer’s protocol. The two Munc13 proteins were then purified using Ni-NTA affinity beads as described before (40-42). Cell pellet was thawed on ice and disrupted with a homogenizer, and then spun in an ultracentrifuge for 30 minutes at ∼142,400 g. 2 mL Qiagen Ni-NTA slurry and 10 uL Benzonase were added into the resultant supernatant, and rotated on orbiting wheel overnight at 4°C. The beads were then washed with buffer containing 25 mM Imidazole. 100 μL of PreScission protease (∼2 mg.mL^-1^) in 1 mL buffer was added to the beads and incubated for 3 hours at room temperature with shaking to remove the 12xHis tag. After cleavage reaction, elusions were collected and gel filtrated using a Superdex 200 column. The protein concentration was typically 1 mg.mL^-1^ as determined by using a Bradford protein assay with Bovine Serum Albumin (BSA) as the standard.

### Protein labeling

The two Munc13-Halo proteins were labeled by incubating the protein with Alexa 488 conjugated with HaloTag® from Promega, as described before (2). The protein was first centrifuged at 14,000 rpm for 20 minutes at 4°C to remove any precipitation. Fluorescence dye was added into the protein solution at dye:protein = 5:1 molar ratio and the mixture were incubated for 30 min at room temperature with gentle rotation. Unreacted dye was removed by passing through the PD MidiTrap G-25 column (GE Healthcare) three times. The labeling efficiencies were about 97%.

### Liposome extrusion

Protein-free liposomes were prepared by extrusion using an Avestin min-extruder (1).

The liposomes used to form bilayer membrane contained 63 mol% DOPC, 25 mol% DOPS, 2 mol% PI(4,5)P2, and 10 mol% DAG. The liposomes used for vesicle capture contained 68 mol% DOPC, 30 mol% DOPS, 2 mol% DOPE-Atto647N. Detailed description was included in the Supporting Information.

Chloroform solutions of the lipid mixtures were dried with a nitrogen stream for 10 min, followed by vacuum drying for 1 hour. The thin lipid films were hydrated with 500 μL buffer containing 50 mM HEPES (pH 7.4), 140 mM KCl, and 10% glycerol. The mixture was vortexed vigorously for 30 minutes at room temperature. The multilamellar liposomes were frozen in liquid nitrogen for 30 sec, and were then thawed in a water bath at 37°C for 30 sec. This cycle was repeated eight times. Small unilamellar liposomes of ∼100 nm were produced by extrusion through 100 nm polycarbonate filters using a Liposofast mini-extruder (Avestin) for 21 times.

### Bilayer Prep and TIRF microscopy

Bilayers were prepared by bursting liposomes on the surface of glass using a glass-bottomed μ-Slide V1^0.5^ chip from ibidi. 2.5 µL MgCl_2_ at 500 mM were added into 122.5 µL buffer containing 50 mM HEPES (pH 7.4), 140 mM KCl, and 10% glycerol, then add 125 µL extruded bilayer liposomes. 60 µL MgCl_2_-liposome solution were loaded into the channel of the ibidi chip and incubate for 40 min. The channel was washed with the same buffer supplemented with 6 mM EDTA, and then with buffer supplemented with 1 mM DTT. 60 µL 10 nM Munc13-halo-AX488 were loaded into the channel and incubate with the bilayer for 60 min. The channel was washed with the buffer supplemented with 1 mM DTT. The vesicle liposomes were diluted 30 times. 60 µL diluted vesicle liposomes were loaded into the channel and incubate for 5 min. The channel was washed with buffer supplemented with 1 mM DTT.

The ibidi chip was then mounted to the stage of a Nikon TIRF microscope. Bilayers, Munc13 particles on bilayers, and vesicles attached to bilayers were respectively imaged with the TIRF microscope using corresponding laser.

## SUPPORTING INFORMATION

Supporting Information includes: Supporting Figures, and Supporting Movies.

## AUTHOR CONTRIBUTIONS

Feng Li, Conceptualization, Reagents, Data curation, Formal analysis, Investigation, Writing— original draft, Writing—review and editing; Venkat Kalyana Sundaram, Alberto T. Gatta, and Jeff Coleman, Reagents, Investigation, Writing—review and editing; Shyam S Krishnakumar, Frederic Pincet, and James E Rothman, Conceptualization, Formal analysis, Supervision, Funding acquisition, Writing—review and editing.

## ACKNOWLEDGEMENTS

This work was supported by National Institute of Health (NIH) grant DK027044 to JER.

## SUPPORTING INFORMATION FOR

### SUPPORTING FIGURES

**Supporting Figure S1.**
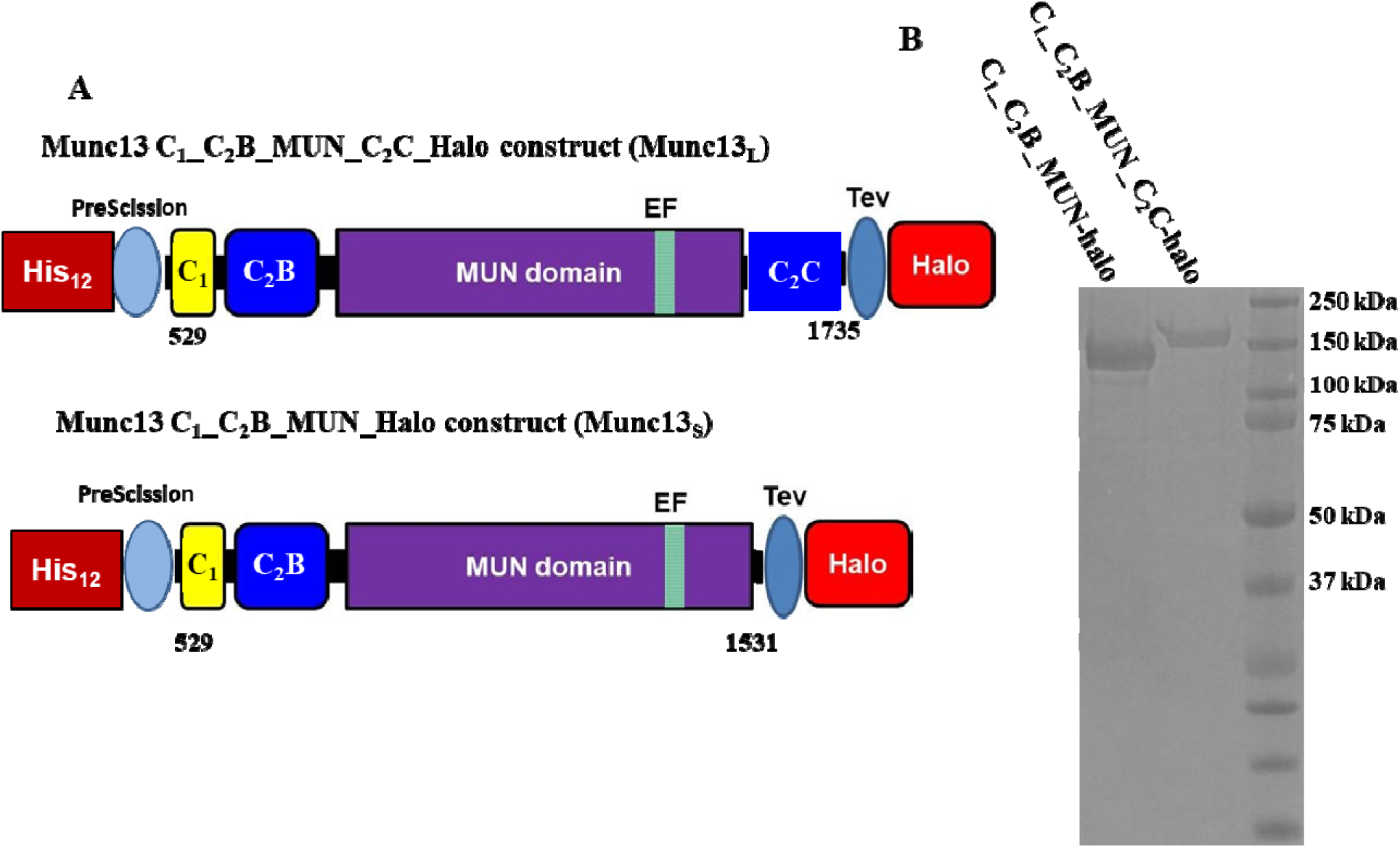
**(A)** Munc13 constructs. (**B**) SDS-PAGE analysis of purified Munc13 proteins.

**Supporting Figure S2.**
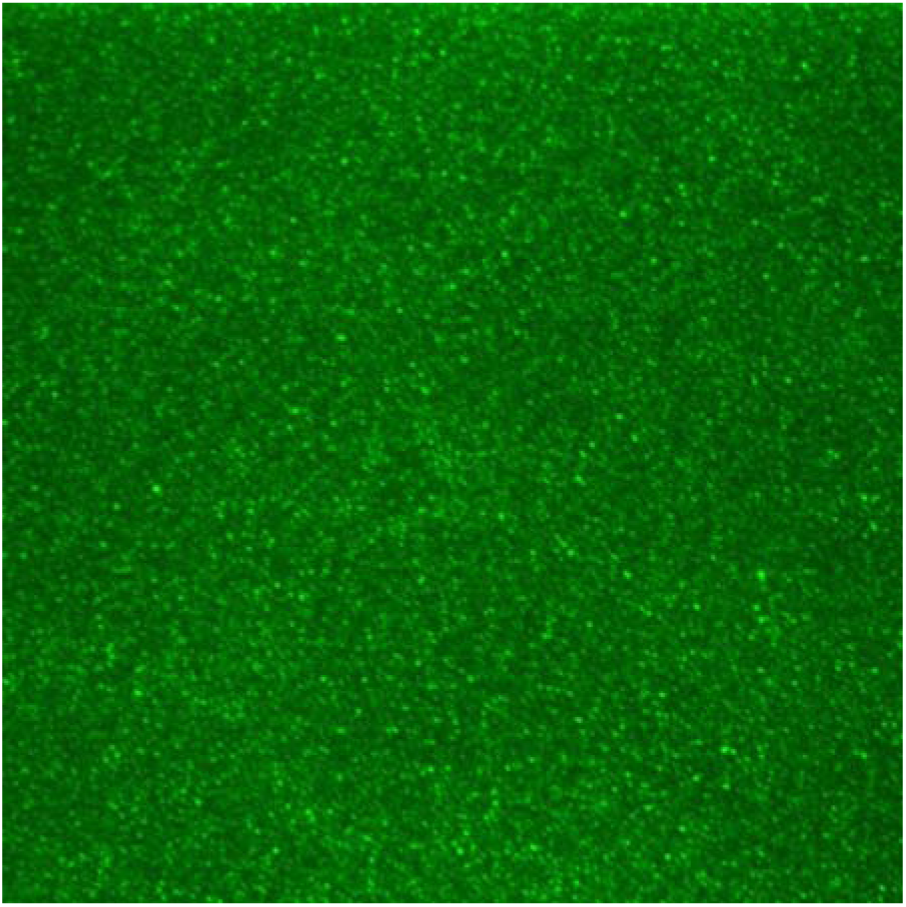
A confocal image of supported bilayer containing DOPC, DOPS, DAG, PIP2 and PE-NBD.

**Supporting Figure S3.**
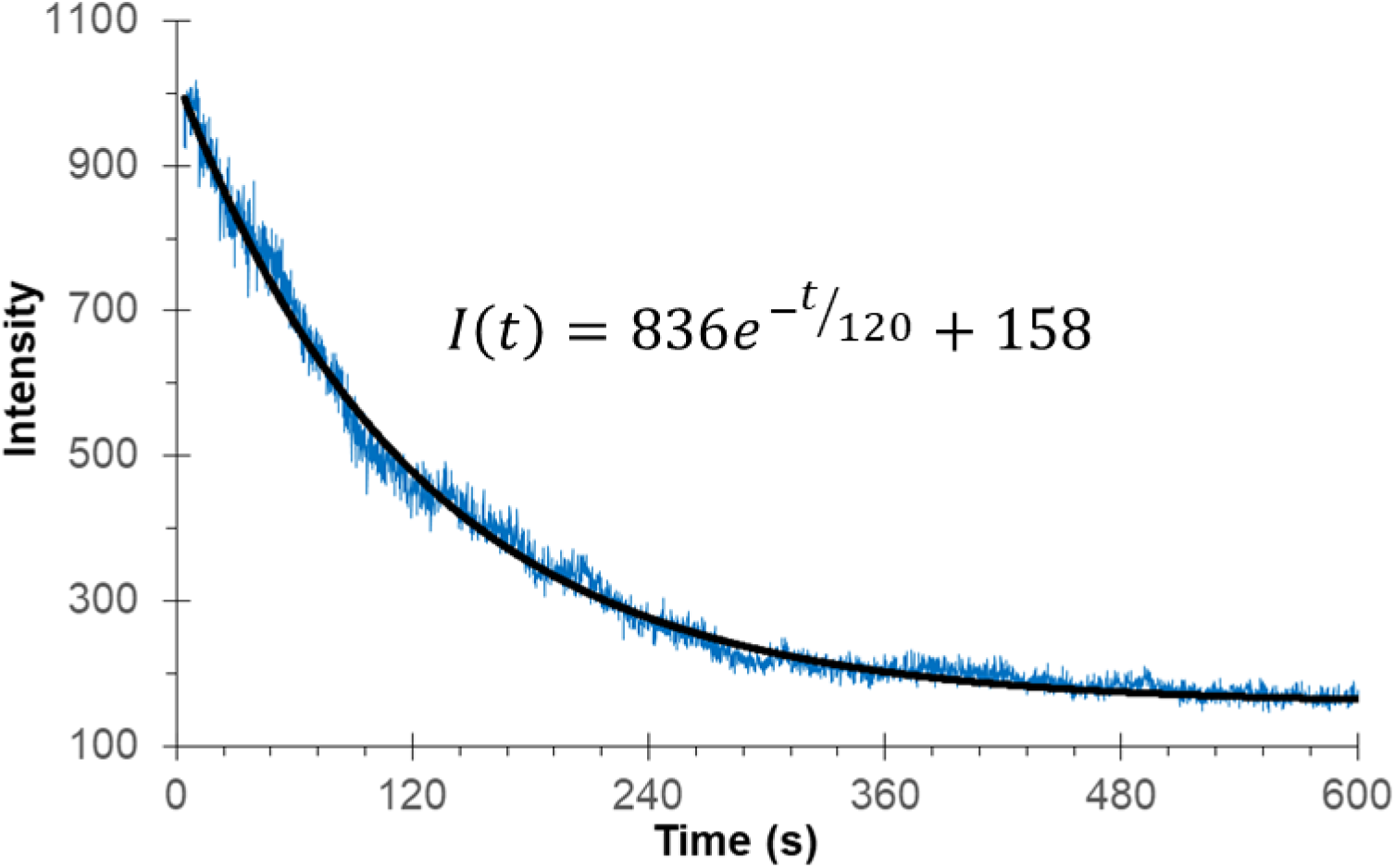
Fitting of the fluorescent bleaching curve *I(t)* vs. *t* to find *I(0)*. The experimental bleaching data (blue curve) was fitted with an exponential equation *I = 836e*^*-t/120*^ *+ 158*. Hence *I(0)* was 836.

### SUPPORTING MOVIES

**Supporting Movie 1. Representative bleaching of Munc13**_**L**_ **clusters formed on lipid bilayer (replay at 8x speed)**.

**Supporting Movie 2. Representative bleaching of clusters by Munc13**_**S**_ Δ**C**^**2**^**C formed on lipid bilayer (replay at 8x speed)**.

